# FASTGenomics: An analytical ecosystem for single-cell RNA sequencing data

**DOI:** 10.1101/272476

**Authors:** Claus J. Scholz, Paweł Biernat, Matthias Becker, Kevin Baßler, Patrick Günther, Jenny Balfer, Henning Dickten, Lars Flöer, Kathrin Heikamp, Philipp Angerer, Mathias Heilig, Ralf Karle, Meike Köhler, Thomas Mazurkiewicz, Martin Mönnighoff, Christian Sauer, Albrecht Schick, Gerhard Schlemm, Roland Weigelt, Martin Winkler, Thomas Ulas, Fabian Theis, Stephan Huthmacher, Christina Kratsch, Joachim L. Schultze

## Abstract

Recent technological advances enable genomics of individual cells, the building blocks of all living organisms. Single cell data characteristics differ from those of bulk data, which led to a plethora of new analytical strategies. However, solutions are only useful for experts and currently, there are no widely accepted gold standards for single cell data analysis. To meet the requirements of analytical flexibility, ease of use and data security, we developed FASTGenomics (https://fastgenomics.org) as a powerful, efficient, versatile, robust, safe and intuitive analytical ecosystem for single-cell transcriptomics.

## Introduction

Recent technological advances increased the resolution of transcriptomics from cell populations (“bulk”) to single cells^1^. While only few cells were assessed in initial projects^2,3^, evolving technologies now allow the analysis of thousands of cells^4–6^, with the largest publicly available dataset currently comprising more than 1.3 million cells7. In contrast to bulk RNA sequencing (RNA-seq), single cell (sc) technologies are much more demanding due to high technical variation with zero-inflation being a major property^8^. As a consequence, a myriad of novel computational approaches and tools have been developed for the different scRNA-seq technologies^9^, but these thriving innovations also constitute a lack of widely accepted gold standards for data analysis. By construction, many of the proposed algorithms and approaches address only certain steps in the analytical scRNA-seq workflow, are adapted to certain scRNA-seq technologies, or cannot be easily combined with other tools, limiting their broad applicability. Notable exceptions are software packages like Monocle^10^, Seurat^11^ and Scanpy^12^, which are well documented, cover big parts of the analysis workflow, and are flexible in their application; nevertheless, due to their command line-based environments, they are still restricting access to scRNA-seq for the broader life and medical sciences community. More user-friendly tools with graphical user interfaces have been introduced, like Granatum^13^, which offers a local installation, or the online tool ASAP^14^ and the commercial solution SeqGeq^15^. In their current versions, they offer popular analysis algorithms, yet are limited in scalability in a multi-user setting, data security, usability of data varying in size over several orders of magnitude, and integration of own analytical concepts. Especially for largest-scale single cell genomics undertakings like the Human Cell Atlas (HCA)^16^, existing tools provide only limited analytical performance due to inefficient resource allocation for exploding memory and computing requirements for datasets in the magnitude of millions of cells, thus underscoring the necessity for a powerful software solution tailored to efficiently handle mega-analyses through distributed computing.

Single cell genomics - with scRNA-seq leading the way - will revolutionize the life and medical sciences^8,17–19^. Here, we postulate that an analytical ecosystem for single cell genomics applications will foster research and development in this field. Such an ecosystem should give computational experts a platform to make their tools available to a broader audience in a user-friendly fashion, allow high-end users to develop individualized workflows, and provide the novice user a computational environment to get acquainted with the special computational requirements for single cell analysis. Furthermore, such an ecosystem should serve as a platform for the community to share public datasets with a broader audience by following the FAIR Guiding Principles for scientific data management and stewardship^20^, provide a scalable infrastructure for projects with large datasets even across numerous institutions, host benchmarking capabilities for newly developed algorithms for the analysis of scRNA-seq data, and even serve as a portal for large international projects such as the HCA^16^. Finally, an analytical ecosystem must implement best practice measures that agree with institutional and governmental data security regulations. To address all these requirements, we have developed FASTGenomics (https://fastgenomics.org) as a powerful, efficient, versatile, robust, safe and intuitive analytical ecosystem for single-cell transcriptomics. Access to the FASTGenomics ecosystem and its functionality is granted for free upon registration to allow unrestricted interaction with the single cell genomics community and especially academia. Furthermore, as suggested by the HCA white paper and guided by representatives of the HCA, the implementation of FASTGenomics as a portal for the HCA is currently on its way.

### App Store in FASTGenomics serves as platform for novel algorithms

At the heart of FASTGenomics is a hybrid app store (**Figure 1A**) optionally composed of public (cloud) and private (local) app repositories hosting algorithms for calculations and data visualization. Novel algorithms can be provided as new apps by the computational biology community (Figure 1B). The well-documented application program interface (API) (**Supplementary Information “Description of the API of FASTGenomics”**) defines data input and output (Figure 1C) and allows seamless integration into the FASTGenomics ecosystem. Apps submitted to the public app repository (https://github.com/fastgenomics) are included in the complete end-user environment (Supplementary Information “Detailed description of end-user experience of the FASTGenomics ecosystem”). Additionally, designing customized workflows integrating custom-made apps is a major feature of FASTGenomics (Figure 1D). Furthermore, the workflow editor allows to adjust parametrization of apps, thus providing a maximum of analytical flexibility. Currently, workflow editing is done via the command line, the next version of the workflow editor is planned to provide an intuitive graphical user interface with functionality to share custom workflows (Supplementary Figure S1).

**Figure 1:**
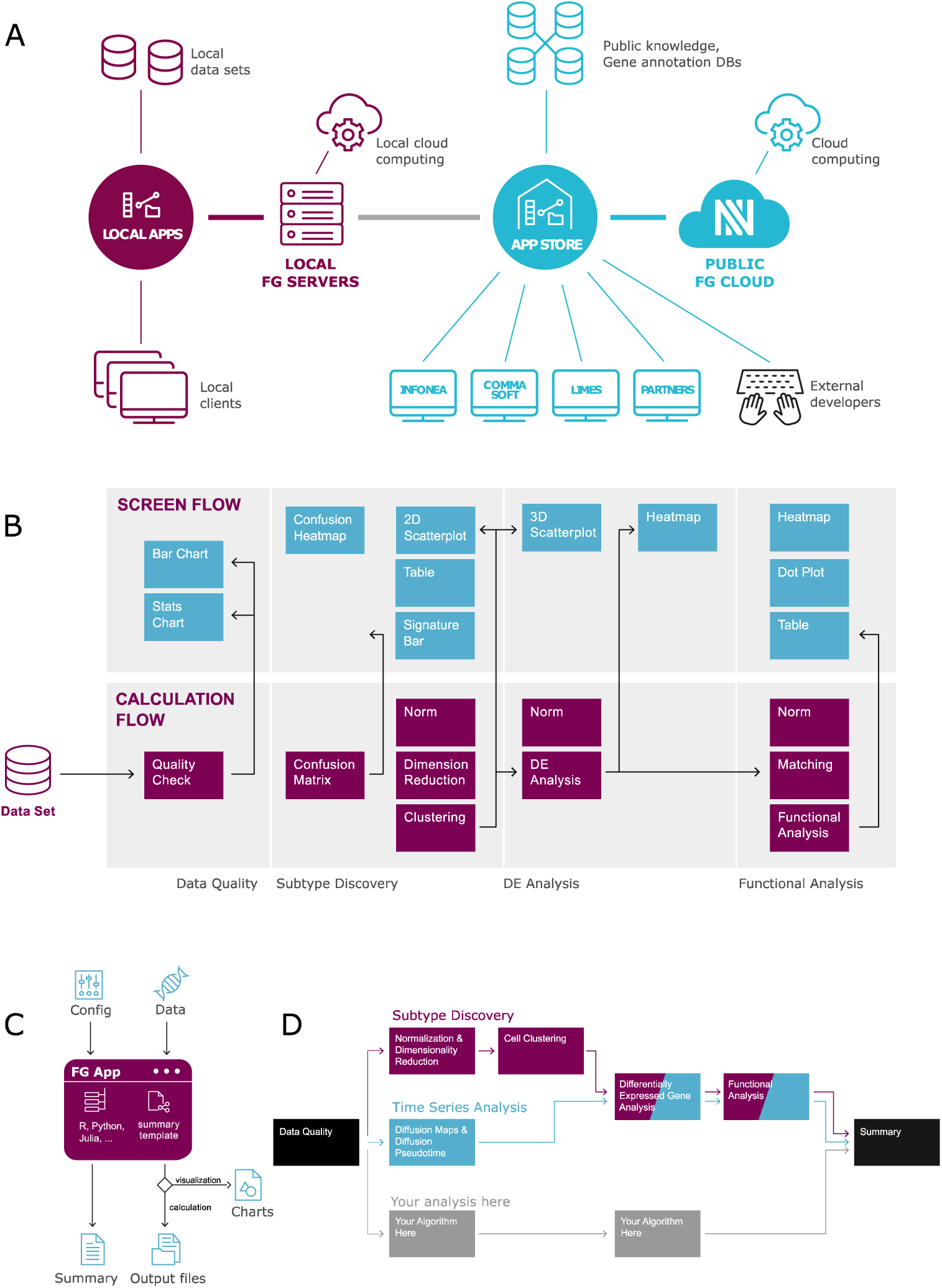
FAST Genomics ecosystem. **(A)** Hybrid app store concept. To provide both the advantages of community access to the FASTGenomics framework as well as the security of a private working environment, FASTGenomics runs in the cloud and can also be installed on premise. The cloud installation allows the usage of public apps and exchange with the global research community, whereas the on-premise installation could run on a local cluster. Additional local app repositories and data storage can be added for private access only. **(B)** Typical structure of a FASTGenomics workflow. All FASTGenomics workflows consist of calculation apps (such as quality checks, data normalization, dimensionality reduction, clustering, …) that take inputs and consecutively produce new results for upstream calculation apps. Selected outputs of the calculation workflow are displayed in the browser with the help of visualization apps in the according visualization workflow. **(C)** Structure of a FASTGenomics app. Apps are Docker containers that interact with the FASTGenomics framework using an interface for data input and a configuration file providing necessary parameters for the analysis. Each FASTGenomics app dynamically generates a summary of the analysis performed by the app that is collected by the FASTGenomics summary service. Depending on app type, different channels are used for results, calculation apps write output to disk, whereas visualization apps send output to theweb browser. The use of the Docker framework enables app developers to implement algorithms in any programming language of choice. A detailed tutorial for the development of calculation and visualization apps as well as sample code can be found at the public FASTGenomics app repository (https://github.com/fastgenomics). (D) Workflow definitions and app concept: workflow definitions are configuration files that describe the calculation and visualization apps used for a specific workflow. User-defined workflows can be added simply by creating new workflow definitions, which may recycle previously defined apps. In particular, apps for the exploration of gene candidate lists with the help of DE analysis and functional annotation are typical candidates for multi-workflow apps. All workflows end with a detailed summary of the analyses performed to ensure maximum transparency and reproducibility.

### Architecture, scalability and data security of the Docker-based hybrid model of FASTGenomics

The FASTGenomics ecosystem has been implemented as a Docker^21^-based cloud solution, which can also be used as a local environment with a community-wide app repository (hybrid design) allowing to share data, apps and workflows, but also information, expertise and knowledge about single cell genomic analyses (**Figure 1A**, for a user perspective, **Supplementary Figure S2** for architectural specifications, for more details see **Supplementary Information “Technical realization of FASTGenomics with Docker-based cloud solution”).** Alternatively, entirely local installations – as they might be required within industry – are also possible. While ensuring standardization and reduced administrative burden, the modular, docker-based hybrid cloud solution of FASTGenomics also provides the necessary scalability to run projects with very large datasets. A dynamic allocation and flexible use of available resources will achieved by leveraging Kubernetes technology in the next release of the platform^22,23^.

In its current version, FASTGenomics is being developed according to EU-GDPR (General Data Protection Regulation) and the German Federal Data Protection Act (“Bundesdatenschutzgesetz”, BDSG), one of the strictest data protection laws in the world. To minimize security issues related to multi-user access to the platform and the use of custom apps, FASTGenomics implements a rigorous multi-layer security concept of data encryption, controlled access and transfer to protect study data (expression tables, sample metadata and analysis results) as well as user data from unauthorized access and manipulation (**Supplementary Figure S3**). A data protection concept has been developed accordingly and will be continuously updated according to legal requirements (**Supplementary Information “Data Security Concept within FASTGenomics”).**

### User-friendly computational environment

Within the FASTGenomics ecosystem, analyses can be initiated and monitored from essentially any web-compatible hardware with a web browser, without requiring extensive computing or memory resources locally. For the end-user following registration, FASTGenomics provides an interface for data upload (**Supplementary Figure S4A, Supplementary Information “Description of data upload via upload Dock in FASTGenomics”)**, starting from count tables and experimental metadata, followed by standardized quality checks, e.g. average molecule counts, gene types, and quantification of batcheffects (Supplementary Figure S4C-E), and two pre-defined data analysis and visualization workflows, ‘Subtype Discovery’ and ‘Pseudo Time Analysis’ (**Figure 1D**). The former includes a neural network approximation of the parametric tSNE^24^ and a 3D visualization of cells with coloring according to cluster assignments, gene expression and metadata (**see also Supplementary Table 1**). Each analysis results in the definition of genes of interest and a functional categorization with the help of external databases, e.g. Gene Ontology (GO^25^). Workflows in FASTGenomics end with a summary, a detailed description of all analysis steps including information about algorithms, software, versions, and parametrizations used as well as input data and results produced (**Figure 1D, Supplementary Figure S5A, S5B, Supplementary Information “Description of Summary of any given analysis”**). The summary is intended to maximize reproducibility and transparency of the analysis, which could be made available e.g. in scientific publications or within documentation required in regulatory environments.

### Platform for sharing datasets for further public exploitation

Another important feature of FASTGenomics is a standardized package for public dataset presentation, which we utilized to present 10 recently published datasets ranging from 482 to 68,579 cells per dataset (**Supplementary Table 2**)^5,26–34^. Available datasets can be connected with standard workflows provided by FASTGenomics, but also with customized apps and workflows as exemplified for a previous MARS-Seq dataset (**Supplementary Figure S6**)^31^. By combining a dataset with the initial analysis, the data can be examined by anybody following the same algorithmic settings as previously reported in the literature. Moreover, this also allows to compare different analysis strategies directly on the same platform. We also performed concordance analyses for selected datasets presented in FASTGenomics (**Figure 2A**) and focus here on a dataset with 3,005 cells published by Zeisel et al.^34^. Using the BACKSPIN clustering algorithm, a total of 9 clusters that were assigned to 7 classes of cell types were previously identified in the dataset, while after our neural network-based dimensionality reduction a subset of 2,375 cells could be assigned to 16 clusters. Thus, the FASTGenomics ‘subtype discovery’ standard workflow revealed a more fine-grained cluster structure than the BACKSPIN algorithm while preserving the co-clustering of functionally closely related cell types. In particular, neuronal and glial cell types were clearly distinguished from each other as well as from vasculature; in more detail, oligodendrocytes and pyramidal neurons were each assigned to one FASTGenomics cluster, while interneurons were clustered to six main classes. Quantitatively this translates to an adjusted mutual information value of 0.75 and median concordance rates of 96.5% for FASTGenomics and 90% for BACKSPIN (**Figure 2AB, Supplementary Information**). Such measures might be also used to estimate specialized analyses settings in previously published datasets. Collectively, the option to freely share previously published large datasets on FASTGenomics allows intuitive and interactive cross-examination, which goes far beyond the current options in scientific publications.

**Figure 2:**
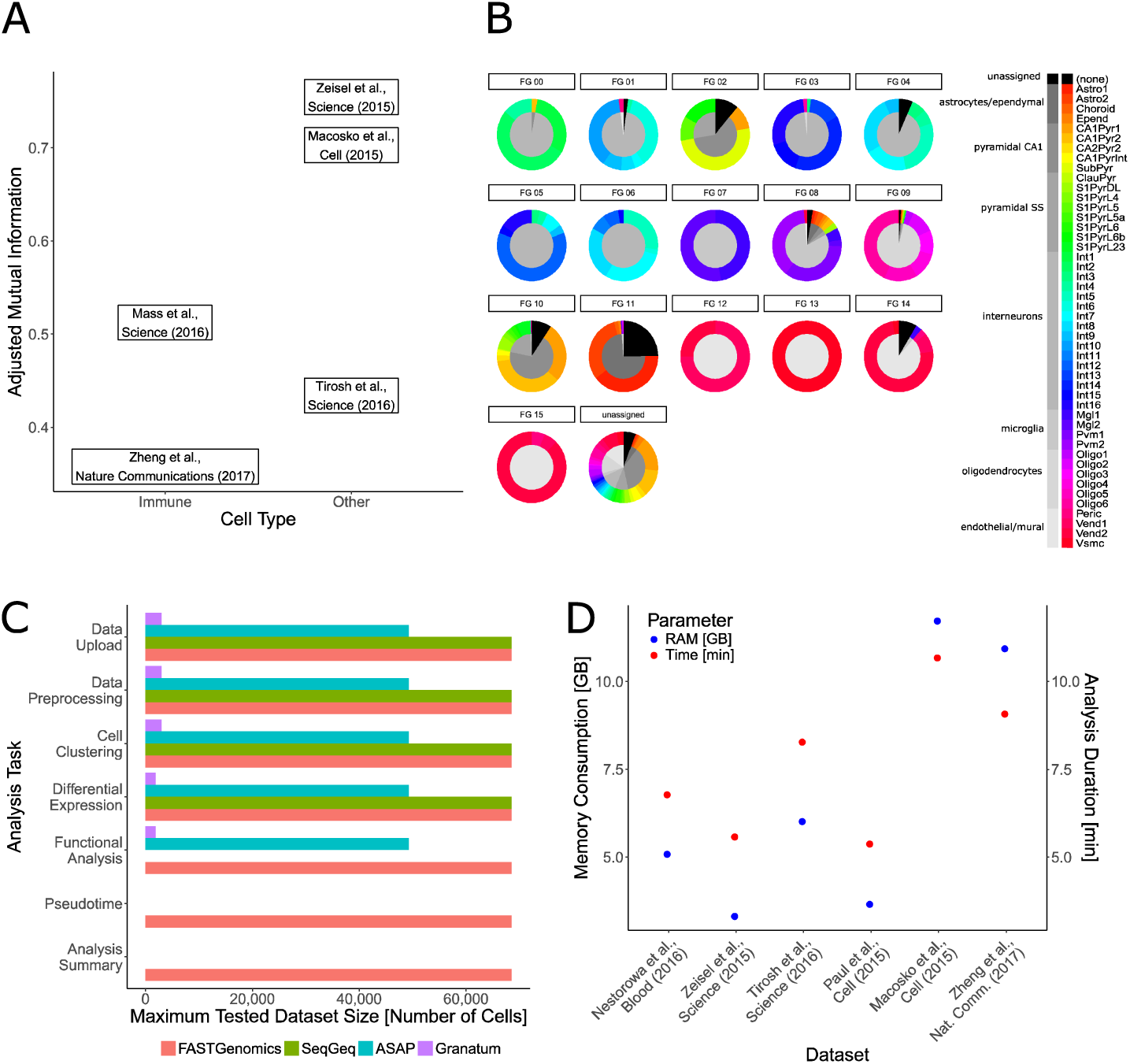
Reproducibility of workflows and performance of FASTGenomics. **(A)** Clustering results of individual cells generated with the standard ‘subtype discovery’ workflow in FASTGenomics were compared to published findings by determination of the adjusted mutual information (AMI). Immune cell datasets^29,31^ displayed a lower degree of concordance than neuronal^34^, cancer^27^ and retinal tissue^5^ datasets, presumably due to the lower RNA content of immune cells^29^ and the lower number of genes expressed^27^. **(B)** FASTGenomics (FG) cluster assignments compared to published cell types^34^. For each FG cluster, the proportion of main cell types (inner circle) and subtypes (outer circle) are shown. The FASTGenomics standard ‘subtype discovery’ workflow clearly distinguished single-cell transcriptomes at higher resolution than main cell types, but with lower resolution than the published subclustering approach. Based on single-cell transcriptomic data, biologically meaningful subclasses were generated by the FASTGenomics ‘subtype discovery’ workflow, classifying neuronal and glial cells, vasculature and immune cell types in distinct units. **(C)** Performance comparison between FASTGenomics and three additional GUI-based platforms for single cell analysis. FASTGenomics (https://fastgenomics.org) was compared to the online tool ASAP (https://asap.epfl.ch/) and local installations of Granatum (http://garmiregroup.org/granatum/app) and SeqGeq (https://www.flowjo.com/solutions/seqgeq) installed on a 64 bit Windows 10 machine with Intel i7 6700K CPU and 32 GB RAM). Comparison was performed in 7 categories (data upload, data preprocessing, cell clustering, differential gene expression, functional analysis, pseudotime analysis, analysis summary). Datasets of various sizes, ranging from 1,920 to 68,579 cells^5,29,32–34^ were used to assess scalability of the platforms. The size of the largest dataset, for which an analysis task could be accomplished is shown for all evaluated pipelines. **(D)** Required resources for analysis of data sets of various sizes^5,29,32–364^. Maximum memory usage (blue dots) and overall analysis runtime (red dots) to complete data normalization, dimensionality reduction and cell clustering are shown depending on the number of cells contained in each analyzed dataset.

### FASTGenomics provides higher flexibility and scalability compared to existing platforms

Next, we intended to compare FASTGenomics to the three currently available GUI-based platforms ASAP^14^, Granatum^13^ and SeqGeq^15^ (for detailed setup **see Supplementary Information “Setup of ASAP, Granatum and SeqGeq for comparison with FASTGenomics”**). We utilized five datasets ranging from 1,920^33^ to 68,579 cells^29^ and compared for data upload, pre-processing cell clustering, differential gene expression analysis, pseudo time analysis and analysis summary. In their default configuration, among the four evaluated tools, only FASTGenomics performed all steps with all datasets (Figure 2C). We furthermore determined the resources needed by FASTGenomics to compute analyses with differeffentdataset sizes (experiment details in “Resource Requirements of a FASTGenomics Analysis Workflow”). Analysis runtime and memory requirements are both strongly correlated and depend on the number of cells analyzed; furthermore, analysis of all datasets across the tested size range is feasible with a contemporary desktop computer (**Figure 2D**).

### Outlook

In upcoming versions of FASTGenomics, datasets, apps and workflows can be shared in private spaces/sections between collaboration partners prior to publishing, thus providing the infrastructure for multi-institutional collaboration projects. Furthermore, import/export apps will be implemented to be fully interoperable with established analysis software tools like Monocle^10^, Scanpy^12^, Scater^35^, Seurat^11^, etc., but also with data repositories like Gene Expression Omnibus (GEO)^36^. Finally, a connection of FASTGenomics to major laboratory information management systems (LIMS) for the import of experimental variables as metadata for new datasets as well as the export of the analysis summary back to the experimenters’ LIMS is currently evaluated and discussed with future users.

## Conclusion

Taken together, FASTGenomics is designed as a secure, flexible, scalable but also standardized platform for single cell RNA-seq data, open to the scientific community. A major feature is to provide highest reproducibility and transparency for single cell data analysis to the whole community. Due to its modular and open structure it could also serve as a platform for community-wide benchmarking for novel algorithms and even serve as one of the tertiary portals planned within the HCA data coordination platform of the Human Cell Atlas^16^. Furthermore, by design, it scales already routinely to more than 5×10^4^ cells per project and prototype apps suggest that scaling to 10^6^ cells is also possible. Moreover, its hybrid design will also allow using FASTGenomics on premise, which might be of interest to clinical research and the pharmaceutical industry.

## Acknowledgements

We would like to thank John Marioni for fruitful interactions concerning the project to implement FASTGenomics as one of the portals for the HCA. This work was supported by a grant from the Federal Ministry for Economic Affairs and Energy (BMWi Project FASTGENOMICS). JLS is a member of the excellence cluster immunosensation. JLS is further supported by the DFG (Sachbeihilfe SCHU 950/9-1; SFB 704, projects A13, Z5; excellence cluster immunosensation).

## Author contributions

CJS wrote the manuscript; CJS, PB, MB, KB, PG, JB, HD, LF, KH, PA, MK and TU designed analyses; CJS, PB, JB, HD, LF, KH, PA, MK, MM, CS, AS and GS implemented apps; HD, MH, RK, TM, MM, CS, AS, GS, RW and MW developed the platform; JB, HD, KH, RK, TM and CS contributed to the manuscript; KH and JB managed the project; SH perceived idea, managed and supervised the project; CK managed and supervised project, wrote the manuscript; JLS, FT discussed and improved the project and the manuscript; JLS perceived idea for the project, managed and supervised the project, wrote and designed the manuscript.

## Competing financial interests

CJS, PB, MB, KB, PG, TU, FT and JLS declare no competing financial interests. PA, JB, HD, LF, KH, MH, RK, MK, TM, MM, CS, AS, GS, MW, SH and CK are paid employees of Comma Soft AG, a commercial company developing the FASTGenomics platform.

