## Supplementary Materials for "FASTGenomics: An analytical ecosystem for single-cell RNA sequencing data"

**Supplementary Information**

**Content:**

- **Description of the API of FASTGenomics**
- **Detailed description of end-user experience of the FASTGenomics ecosystem**
- **Technical realization of FASTGenomics with Docker-based cloud solution**
- **Logical implementation built on the Docker-based cloud solution**
- **Adding third party apps to the Docker-based cloud solution**
- **Data Security Concept within FASTGenomics**
- **Description of data upload to FASTGenomics**
- **Description of Summary of any given analysis**
- **Concordance rate analysis of FASTGenomics Analyses with published results**
- **Neural network-based dimensionality reduction and clustering**
- **Methods applied to determine concordance**
- **Results of concordance rate analysis**
- **Setup of ASAP, Granatum and SeqGeq for comparison with FASTGenomics**
- **Supplementary Figures 1 to 6 (including legends)**
- **Supplementary Tables 1 and 2**

**Description of the API of FASTGenomics**

The FASTGenomics ecosystem for single-cell analyses allows integrating algorithms from third parties. For proper integration into the FASTGenomics pipeline, implementations must comply with the FASTGenomics application-programming interface (API). This requires that apps are provided as Docker containers of defined structure, with version information provided in Docker tags. The root directory must include a `Dockerfile`, the script to prepare and invoke app components. A `manifest.json` file that contains app descriptions, input, output and parameter definitions. The `sample_data` folder containing files required for integrity tests during app validation. The `readme.md` file with the app documentation and the source code providing the app functionality. Further optional or best practice components of a generic FASTGenomics app include the `docker-compose.yml` file with information how to build and start the Docker container and providing input/output directories. A `requirements.txt` detailing dependencies for proper app functioning. Markdown-formatted template file(s) for the summary output that is dynamically filled with information generated during the app run. FASTGenomics distinguishes between two types of apps that interact with different parts of the pipeline, calculation apps are handled by the workflow engine, while visualization apps are managed by the workflow client. Both app types use defined mountpoints for data (`/fastgenomics/data`, read-only) and analysis configuration file input (`/fastgenomics/config`, read-only). Calculation apps write results (`/fastgenomics/output`, read/write) and a summary (`/fastgenomics/summary`, read/write) to disk, whereas visualization apps send output to the web browser via port 8000. Apps developed to comply with the FASTGenomics API can be published in the public FASTGenomics app repository (https://github.com/fastgenomics) and will be made available in the FASTGenomics Docker registry. A detailed tutorial for the development of calculation and visualization apps as well as sample code can be found at https://github.com/fastgenomics.

**Detailed description of end-user experience of the FASTGenomics ecosystem**

FASTGenomics (https://fastgenomics.org) allows several distinct levels of usage and access. An anonymous access allows users interested in FASTGenomics to see results from pre-calculated analyses on a selection of publicly available datasets. Full access to all functionality requires a free user registration, which enables usage of a greater set of experiments from public repositories, the possibility to integrate own data sets, as well as the availability of the full range of analysis tools. The natural first step of a new project is to upload single-cell expression data to FASTGenomics after which the data may be checked for quality by exclusion of cells with too few expressed genes, and exclusion of genes expressed in too few cells (**Supplementary Figure S4E**). The according calculation app can be incorporated into the workflows to perform this task (**Supplementary Table 1A**). This step is followed by the data quality screen showing general statistics about the data, such as average molecule counts and quantification of batch effects. For data analysis, FASTGenomics offers two pre-defined alternatives: subtype discovery and time series analysis (**Figure 1D**). Furthermore, the workflow editor allows creating custom analytical scenarios that can involve any app available in the FASTGenomics app store. In both pre-defined workflows, an overview of the dataset detailing aspects of data quality (e.g. summary statistics on expression values, presence of putative batch effects, etc.) is given in the first screen (**Supplementary Figure S4B-D**). The subtype discovery workflow proceeds with data normalization and dimensionality reduction, followed by clustering of cells (**Supplementary Figure S1C,D**). This cluster projection is displayed in an aquarium plot, with the first two dimensions corresponding to coordinates determined in our parametric t-SNE approach and the third dimension representing the cluster assignment confidence (with high-confidence cluster assignments "swimming" on top and low-confidence assigned cells sinking to the ground). The next step detects differentially expressed genes between clusters and display these in a heatmap (data not shown). In the pseudotime workflow, diffusion maps are generated to order cellular transcriptomes along pseudotemporal axes. This workflow also determines genes responsible for branches in the trajectory. Both, the subtype discovery as well as the pseudotime workflow conclude the analytic sequence with the functional characterization of signature genes using gene ontology enrichments. At the end of all workflows, a detailed summary is dynamically generated during runtime and displayed to the user (**Supplementary Figure S5B**).

#### Technical realization of FASTGenomics with Docker-based cloud solution

The publicly accessible instance of the FASTGenomics ecosystem (https://fastgenomics.org) is installed in the Microsoft Azure cloud hosted by server infrastructure located in Western Europe. Currently, FASTGenomics services run on a Standard D8s v3 system (8 vCPUs based on the 2.3 GHz Intel XEON ® E5-2673 v4 processor and 32 GB RAM). However, other options can be envisioned since the system is designed to allow hybrid computing by integrating local and public cloud installations.

The FASTGenomics ecosystem itself is based on Docker infrastructure (currently using version 17.06.0-ce, build 02c1d87), all pipeline components as well as calculation and visualization apps are packaged in Docker containers. Internal pipeline components and apps are deployed to different Docker registries, the former can only be accessed by the runtime environment and FASTGenomics administrators, while the latter is also accessible to the community to allow contribution of apps.

The FASTGenomics ecosystem consists of several major components, each with its own responsibility as shown in **Supplementary Figure 2**. The user directly interacts with the `FASTGenomics Client` via the web browser. This serves to display the FASTGenomics website, where users can login, upload and select datasets, choose analysis workflows and access other specialized services. A calculation engine consisting of the `Workflow Engine`, the `Task Dispatcher` and the `Container Service` manages the application of an analysis workflow on a selected dataset. The `Workflow Engine` uses the `Container Service` to start and stop calculation apps that perform individual analysis steps. The latter reports the calculation status to the `Task Dispatcher`, that initializes (creates a unique ID) and finalizes analysis instances upon requests by the `Workflow Engine`. The `Workflow Client` shows a screen flow in the web browser to display results from analyzes; this also makes use of the `Container Service` to start and stop visualization apps. Both components access the `Data Store`, which organizes data management for an analysis. Finally, the data upload system consists of the `Upload Client`, which takes care of data transfer from the user's system to the FASTGenomics servers, as well as the `Packaging Service` integrating the uploaded data into the FASTGenomics system. Overall, connection to the user is secured by an `OpenID Connect` component to ensure that only validated users can access the application.

##### Logical implementation built on the Docker-based cloud solution

Each FASTGenomics project consists of one or more analyses and the data set that is to be analyzed. An analysis in turn can be broken down into the algorithms that calculate results and the visualizations that display these results (**Supplementary Figure S2**). In FASTGenomics, the former are called calculation apps and are combined into workflows, while the latter are called visualization apps and encapsulated into screen flows.

##### Adding third party apps to the Docker-based cloud solution

The scientific community may develop individual apps for FASTGenomics, which are wrapped up in Docker images. A provider of apps is authorized to push Docker images to the public FASTGenomics Docker registry where the images are retrieved on demand by the system. Apps can be calculations producing results or visualizations displaying results. To use a custom app in a FASTGenomics analysis, workflow or screen flow definitions are adjusted. In the current online release, two pre-defined analysis workflows are integrated. Adding custom apps and analyses is a feature of FASTGenomics that is predicted to be active and under continuous development.

##### Data Security Concept within FASTGenomics

The FASTGenomics ecosystem as well as any other web-accessible multi-user platform storing and analyzing sensitive data (e.g. unpublished experimental data, clinically relevant metadata, user data) are subject to tight regulations for data security. As maintainer of an online platform, FASTGenomics needs to adhere to the law including the German Federal Data Protection Act ("Bundesdatenschutzgesetz", BDSG[1]) and of May 2018 the European General Data Protection Regulation (GDPR) (Regulation (EU) 2016/679[2]). These regulations cover diverse aspects of data management and safety.

In order to respond to these regulations, FASTGenomics is continuously working on a security concept defining the essential, recommended and desirable security features of a single cell analysis platform. This is an ongoing process to account for new developments and planned future components and functionality. At the current state, FASTGenomics has the following security features implemented:

- All data in FASTGenomics are stored on encrypted data volumes.
- To ensure safe network topology, both the external and the internal communication between the components is encrypted by HTTPS.

- Authentication is achieved by an OpenID Connect Provider. After registration, users are required to confirm their identity via mail to avoid platform misuse by bots.
- While accessing the platform and user data therein, access is again regulated via authorization checking done by each involved application. This feature manages access rights of each user, for example when retrieving information from the data module. Here, the platform ensures that only eligible private and public data sets are visible for the current user (**Supplementary Figure S3**).
- Finally, several security features address the setup of the Docker container making up the FASTGenomics infrastructure. No Docker container is allowed to have root access. Containers that communicate with external components enforce authenticated users and only communicate using HTTPS. The export of a port is limited to this container group.
- Implementation of national legal requirements for intellectual property with respect to software development (app development).
- Definition and monitoring of organizational best practices for all processes involving data handling, resource access allocation and platform administration.
- Definition and implementation of best practices for internet access of apps.

In addition to these security features, the FASTGenomics security concept addresses further actions for risk minimization and data protection as features planned for future development. These include:

- implementation of a framework for error logging, data access, and data manipulation
- rules that provide manipulation security of data and apps
- definition of best practices in the context of software development in general and the use of container frameworks like Docker in particular, e.g. managing resource access of apps
- definition and implementation of rules for computing resource access of apps
- software support for complete data removal upon user request
- definition and implementation of best practices for validation of usage statistics and application of web tracking software (e.g. Google Analytics)
- definition and implementation of best practices for anonymous access to suitable resources
- definition and implementation of best practices for data publication

Apart from ensuring data security, FASTGenomics aims to provide a good framework for the reproducibility of analyses and the sharing of data and knowledge. Such aspects are increasingly discussed in the scientific world, and driven by concepts like FAIR aiming to facilitate research and knowledge discovery by Findable, Accessible, Interoperable, and Re-usable data[3]. Therefore, the security concept is continuously extended to suggest best practices for data sharing, data publication, community features on the platform, e.g. user forums, use of the summary feature, or use of social media within the platform.

**Description of data upload to FASTGenomics**

FASTGenomics allows registered users to upload own data sets in the data module for further analysis. In the data upload window, the user is asked to provide the expression file in sparse format, the NCBI taxonomy ID of the organism and a title for the later appearance in the dataset item list. Once the user has provided the information, the uploaded data is transformed to our internal FASTGenomics Data Package Format. A Python script checks the NCBI Entrez IDs and those gene IDs that cannot be mapped uniquely. The script automatically documents the removed genes. Once finished, the data is provided to the user in the data module. Thereby, the data set is by default only visible to the person who has uploaded it.

In addition, we also provide the possibility to directly generate the FASTGenomics Data Package Format. A documentation including an R-based tutorial for dataset preparation can be found at https://github.com/FASTGenomics/FASTGenomics_Data_Package_Format. Here, the user can add further information to the data set like metadata for the cells and genes and a dataset description including a short abstract, and contact information. This FASTGenomics Data Package can then be uploaded to the data module and allows fast access to the FASTGenomics functionalities.

#### Description of Summary of any given analysis

The analysis summary report gives a detailed overview of all steps executed and results produced in an analysis workflow to facilitate understanding and to ensure reproducibility (**Supplementary Figure S5**). Workflows describe the sequence of analysis steps performed to get from raw data to an analysis result. Such workflows are not necessarily linear and may contain several branches (e.g. when one app depends on input from several other apps executed before), i.e. a workflow is a directed acyclic graph. Each node (i.e. a versioned calculation or visualization app) in this graph is described in the workflow definition, including information on the analysis context within the workflow. Furthermore, each app provides text passages containing information about applied methods and offers links to access generated interim results during runtime. For report generation, the `summary_visualization` app recursively resolves the app dependencies (required input data and necessary analysis steps to generate this) from leaf to the root and dynamically assembles information gathered from the nodes into the final analysis report. Apps connecting the analysis summary to laboratory information management systems (LIMS) will be developed in the near future and included into the FASTGenomics analytical ecosystem.

#### Concordance rate analysis of FASTGenomics Analyses with published results

Analysis of single-cell RNA-seq data is a complex multistep procedure with many methods available for individual tasks, however with no gold standard being defined. Published experiments thus typically present analysis strategies that are highly specific for the respective underlying dataset. Accordingly, comparison of analysis strategies can be a daunting task. Here, we applied the FASTGenomics subtype discovery workflow for the analysis of a selection of published single-cell RNA-seq datasets generated with different technologies and of various dataset sizes. One common task in single-cell RNA-seq analysis is the definition of cell clusters to define sub-populations in complex mixtures of cells, with a definition of characteristic gene expression signatures and their functional characterization being typical downstream applications that crucially depend on the cluster assignment of cells. We therefore quantitatively compared FASTGenomics single-cell cluster assignments based on a neural network-based dimensionality reduction algorithm (see description below) to previously published clustering results for a selection of single-cell RNA-seq datasets (**Supplementary Table 2**)[4–9].

**Neural network-based dimensionality reduction and clustering**

The standard subtype discovery workflow in FASTGenomics consists of three calculation apps that reduce the input dimensionality and group samples based on their similarity as seen in the gene expression profile. The first calculation app normalizes the data using the term-frequency times inverse-document-frequency (TF-IDF)[10]. This scheme replaces the gene expression in each sample with a number proportional to the expression amplitude in this sample multiplied by the inverse number of samples in which the gene is observed. This amplifies genes which are specific for a given subpopulation and dampens the effect of genes which are present in most samples. Since the non-linear dimensionality reduction needs a dense matrix with intermediate dimensionality, the sparse, normalized expression table is compressed to 32 dimensions using truncated singular value decomposition implemented in a second calculation app. In the third step, a calculation app uses a neural network to approximate a parametric t-SNE embedding[11]. This step projects the intermediate 32-dimensional data onto a two-dimensional space. The neural network approximates the t-SNE optimization problem by learning a projection that minimizes the t-SNE loss function and allows iterative training on batches. By default, the app uses batches consisting of 512 samples and calculates the joint probabilities of samples in the higher dimensional space (by default 32 dimensions from the truncated singular value decomposition) and the target two-dimensional space. Then, the network minimizes the Kullback-Leibler (KL) divergence between input probability and output probability similar to the original t-SNE algorithm. Finally, clustering of the cells is performed using the HDBSCAN algorithm.

**Methods applied to determine concordance**

***Dataset pre-processing and clustering***

For cluster determination we downloaded count tables derived from previously published scRNA-seq datasets (**Supplementary Table 2**) and used them as they were provided within the public repository. After upload into FASTGenomics we used the pre-installed workflow 'subtype discovery' for the identification of clusters within the dataset using the above described neural network-based dimensionality reduction algorithm. Cluster assignments for individual cells were requested from the corresponding authors of individual datasets and compared to those obtained by the FASTGenomics workflow.

***Concordance rate calculation***

From individual cells' published cluster assignments as well as clusterings produced with the FASTGenomics subtype discovery workflow, a contingency matrix $M$ of cell counts per cluster pairs was generated. To provide a quantitative measure for the concordance of clustering results obtained from the two methods, we calculated the concordance rate $C$ for each cluster produced with a specific method as follows:

$$C(M_i) = \frac{max(M_{i,} \cdot)}{\sum_j M_{i,j}} \times 100$$

To provide an overall summary statistic how well one clustering method captures results from the other method, the median concordance rate was calculated. For each pair of clustering methods, two median concordance rates (one for each method) can be calculated.

***Adjusted mutual information***

To quantify the overall clustering concordance between published and the FASTGenomics analysis, we use the adjusted mutual information (AMI), which ranges between 0 (two clusterings show only random overlap) and 1 (overlap between two clusterings is not due to chance)[7].

**Results of concordance rate analysis**

To provide an unbiased analysis of the publicly provided dataset we did not adjust the number of cells when uploading the data to the FASTGenomics portal. As shown in **Supplementary Table 2**, the number of cells reported in the respective publications and the number of cells publicly available was not always identical. Due to the inclusion of different sets of cells into the analysis and the differences in analysis settings, we are aware that the number of clusters between the FASTGenomics analysis and previously published results can vary. Nevertheless, if publicly available datasets are to be used by a broader community, we postulated that the use of the complete datasets provided will be the default usage of such data resources.

We performed a quantitative comparison between previously reported cluster structures and clusters determined in an unbiased fashion by the FASTGenomics 'subtype discovery' pipeline for a selection of single-cell RNA-seq datasets (**Figure 2A**). We observed variation in the AMI values determined for the selected datasets and hypothesize that apart from technical influences like sparsity of the expression matrix and read coverage per cell, also biological aspects impact the clarity of cellular subtype discovery. We found that AMI values were generally lower in immune cell datasets compared to those from other cell types (cerebral and cancer cells as well as retinal tissue), presumably due to the lower RNA content of immune cells and the lower number of genes expressed therein[8,9].

A more detailed analysis of the cluster structure was performed for 3,005 single-cell transcriptomes derived from the murine primary somatosensory cortex (S1) and the hippocampal CA1 region[4], which was previously divided into 9 main clusters and 47 subclasses. When applying the FASTGenomics 'subtype discovery' pipeline, we identified 16 clusters. Of the 3,005 cells analyzed in the published study[4], 630 cells (20.1%) could not be assigned to any of the clusters due to limited assignment confidence resulting from almost equidistant positioning between cell clusters in the tSNE space. However, the median concordance rates for the previously determined 9 main clusters and the 16 newly defined ones were as high as 96.5%

for FASTGenomics and 90% for BACKSPIN, arguing for a high degree of concordance for a large fraction of clusters and cells. Likewise, the AMI for both clustering results was high ($AMI_{Zeisel\ et\ al.,\ Science\ (2015)} = 0.75$, **Figure 2A**). We further compared the 16 FASTGenomics clusters to the 7 cell classes corresponding to the 9 published main clusters as well as to the 47 cell subtypes (**Figure 2B**). Here, all oligodendrocytes classes and all S1 and CA1 pyramidal neurons were each captured by one FASTGenomics cluster, while interneurons were mainly represented in six distinct clusters by FASTGenomics. Thus, the standard subtype discovery workflow revealed an intermediate resolution between the overall and the fine-grained published analysis without generating contradictions to existing knowledge.

#### Setup of ASAP, Granatum and SeqGeq for comparison with FASTGenomics

To evaluate the analytical capabilities of the FASTGenomics analysis pipeline, we defined a set of analytical tasks and checked the performance on published single-cell analysis pipelines featuring a graphical user interface, ASAP[12], Granatum[13] and SeqGeq[14]. ASAP was evaluated using the publicly accessible online instance at https://asap.epfl.ch. Granatum required the local installation of VirtualBox version 5.1.26 (Oracle) and the import of the Granatum appliance version 1.1_2 obtained from http://garmiregroup.org/granatum/app. SeqGeq version 1.3 was obtained from https://www.flowjo.com/solutions/seqgeq and installed locally in the default configuration. The performance of Granatum and SeqGeq was evaluated on a 64-bit Windows 10 machine with Intel i7 6700K CPU and 32 GB RAM.

#### Resource Requirements of a FASTGenomics Analysis Workflow

The memory requirements and the computing time to complete an analysis of a dataset of defined size were chosen to describe the performance of the FASTGenomics pipeline. These parameters were determined in a single-user setting for datasets consisting of 1,920 to 68,579 cells[4,5,7–9,15]; analysis tasks evaluated for the performance measurements were data normalization, dimensionality reduction and cell clustering, because the effort needed for detection of differentially expressed genes and their functional analysis depend on the number of clusters found in the single-cell dataset. Computing times for individual analysis steps were extracted from the Docker log generated by the FASTGenomics Task Dispatcher and summed up for all steps in the analysis workflow. Memory requirements were determined with a batch script running in the background during the calculations that executes `docker stats` and `docker ps` in intervals of two seconds logging memory consumption of individual containers. During the runtime of each container, the maximum memory requirement was used for further evaluation. Resource requirements were determined with the publicly accessible instance of FASTGenomics, which is currently installed on a Standard D8s v3 system (8 vCPUs based on the 2.3 GHz Intel XEON ® E5-2673 v4 processor and 32 GB RAM).

**Supplementary Figures**

### Figure S1

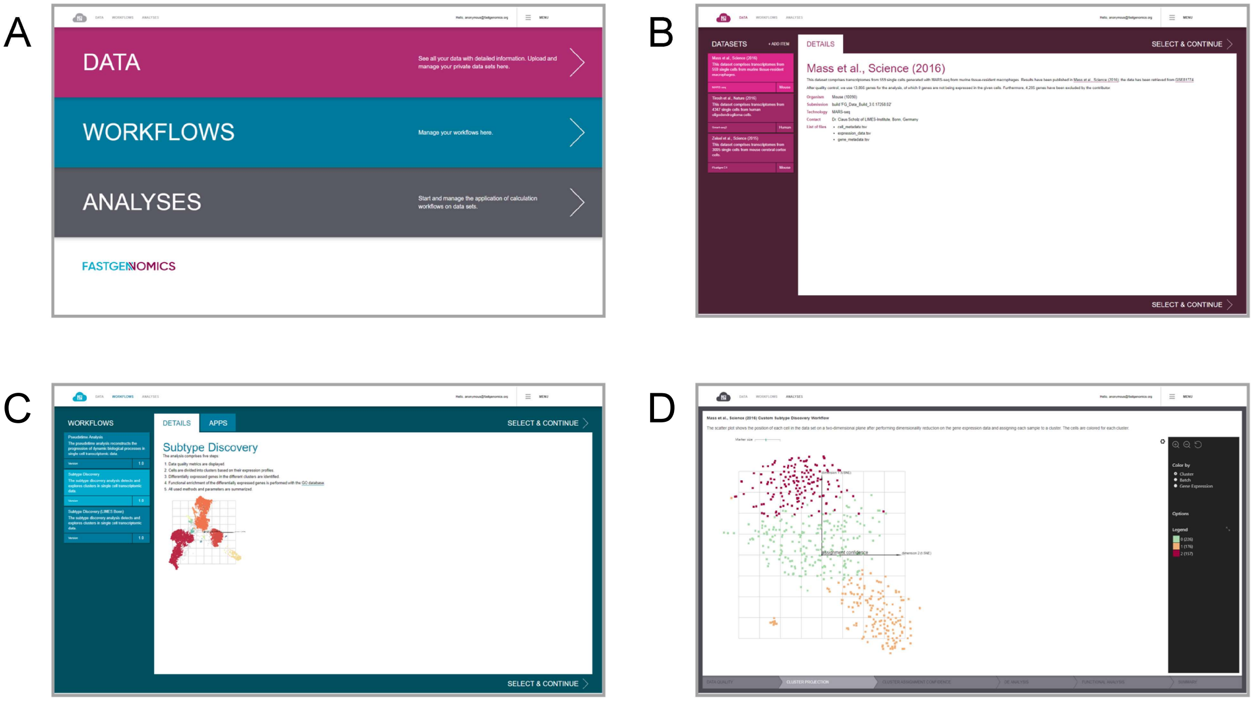


**Supplementary Figure 1: Graphical User Interface of the FASTGenomics analysis ecosystem. (A)** The main menu. **(B)** Overview of accessible datasets. **(C)** Workflow selection page. **(D)** Analysis result visualization.

#### Figure S2

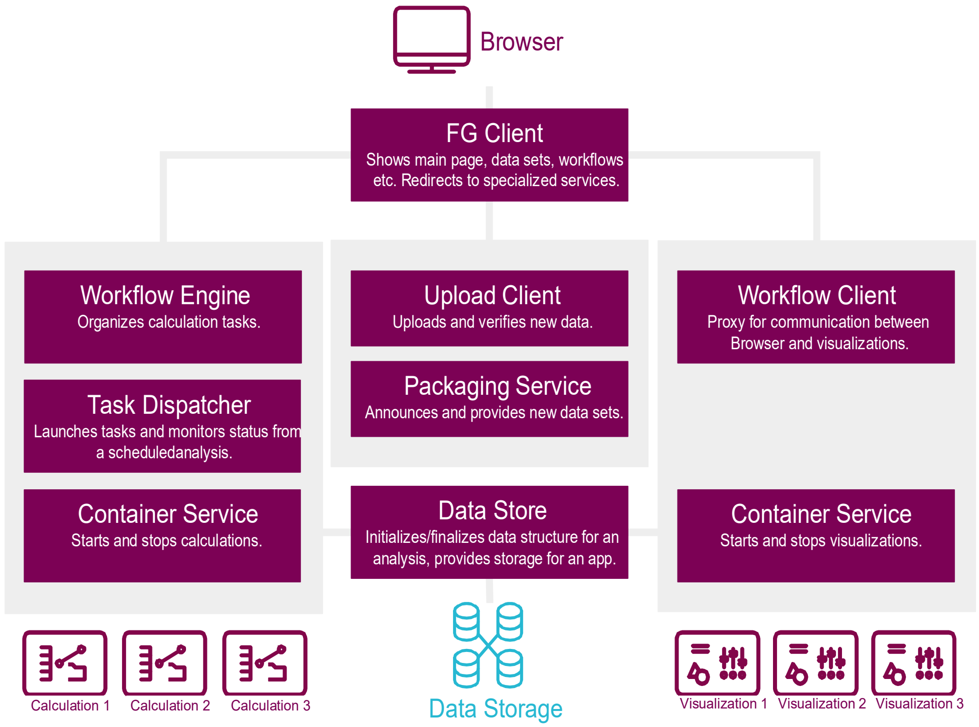


**Supplementary Figure 2: Technical Representation of the FASTGenomics Architecture.** A web browser allows the interaction with the FASTGenomics Client, which internally manages the upload dock (middle branch), the calculation engine (left branch) and the screenflow engine (right branch).

Figure S3

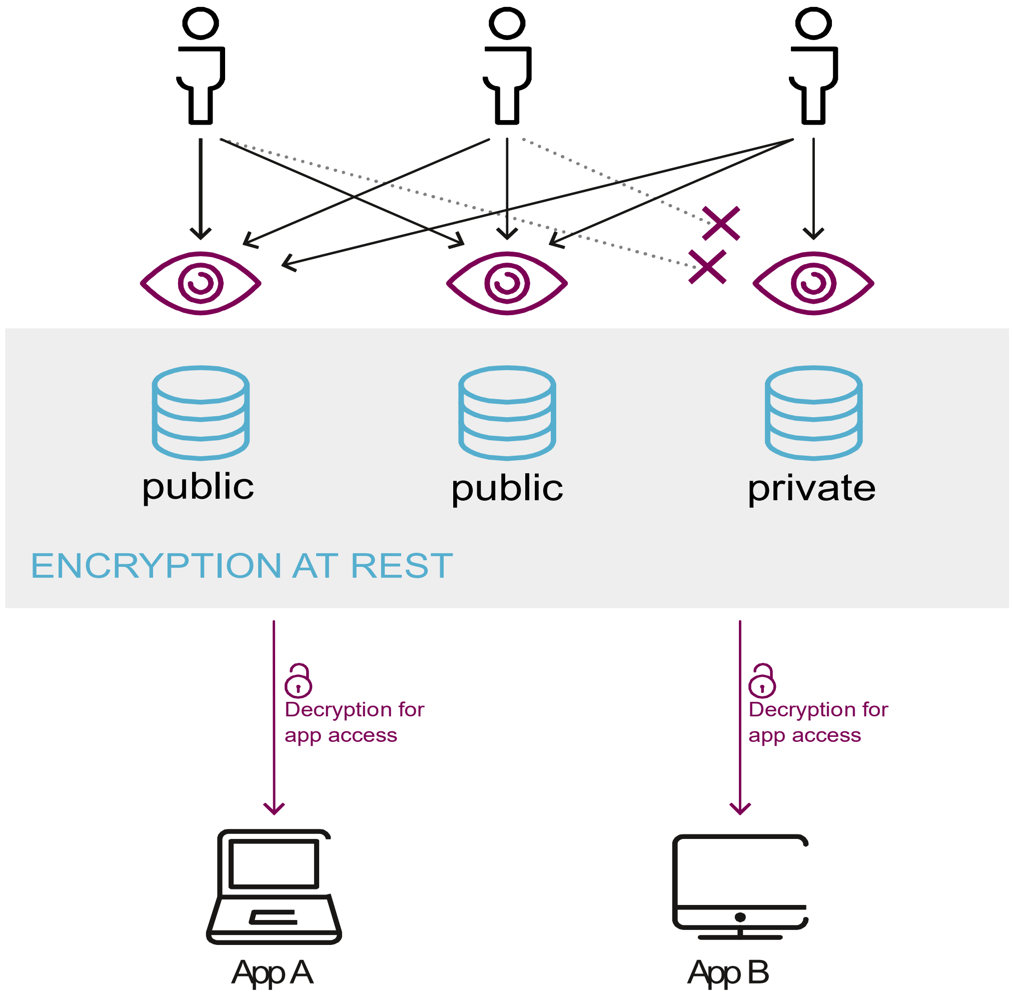


**Supplementary Figure 3: Data Access.** All data is stored on encrypted storage devices. Users can access public datasets and those uploaded by the users, but nor those from other users. Data is encrypted for calculation and visualization apps.

### Figure S4

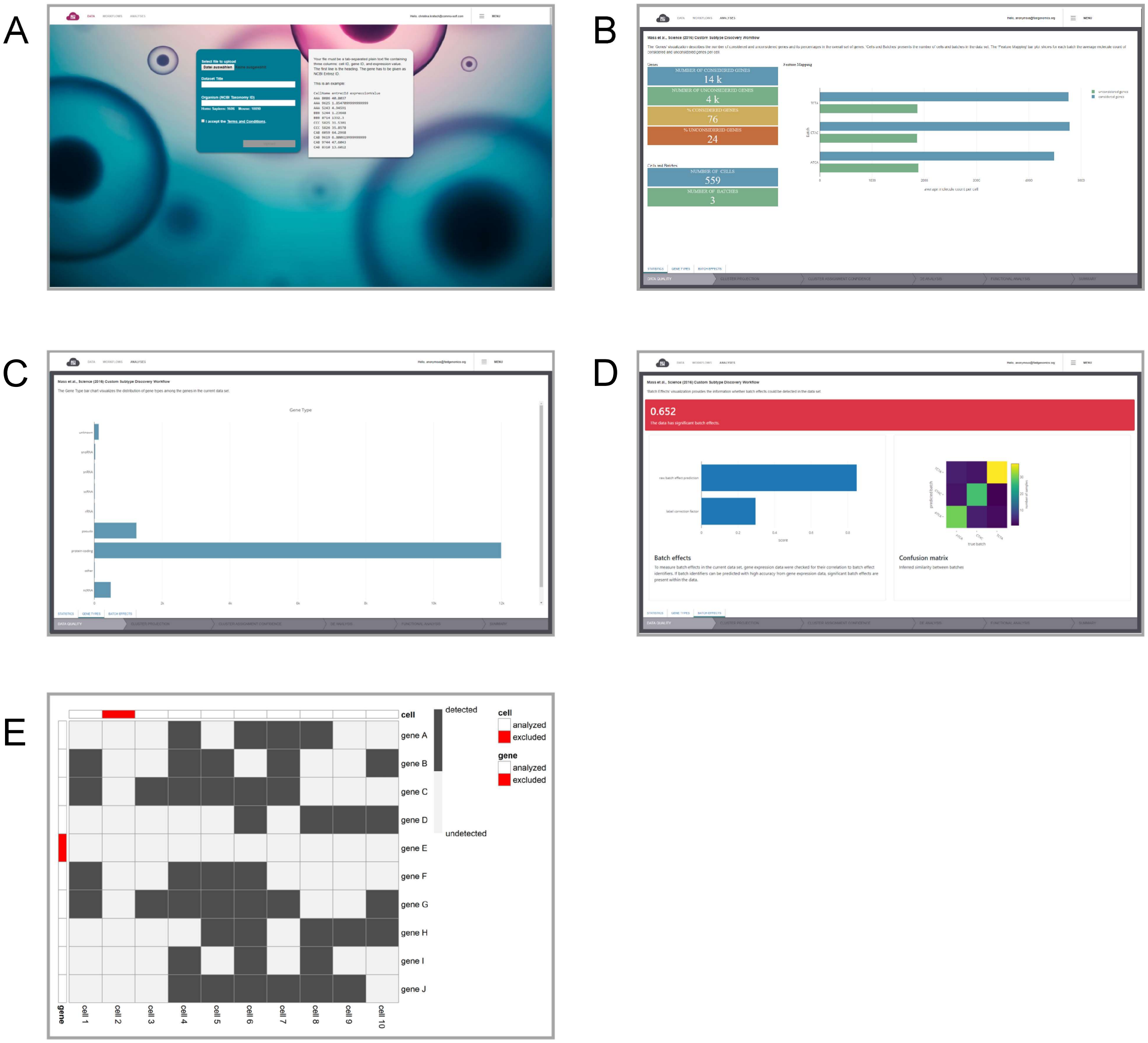


**Supplementary Figure 4: Data Quality. (A)** Data upload in FASTGenomics. **(B)** Data quality check concept. Based on the overall number of genes detected in the single-cell transcriptomics dataset, a lower threshold for the number of genes per cell is dynamically generated; cells expressing fewer genes are excluded from analysis. Furthermore, genes expressed in less than a predefined proportion of cells are removed from the dataset. **(C-E) Quality check screens in FASTGenomics**, screenshots illustrating **(C)** average molecule counts, **(D)** gene types, **(E)** quantification of batch effects.

### Figure S5

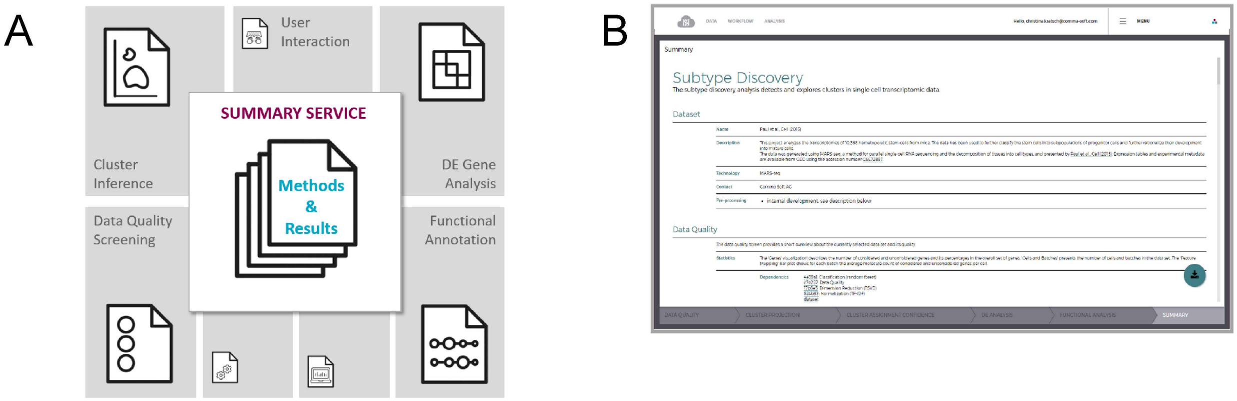


**Supplementary Figure 5: Analysis Summary. (A)** Schematic overview of the summary report-generating app. Information about analyzed data, applied workflows with invoked apps including version information and parametrization are recursively resolved and compiled during run-time to generate the summary report. **(B)** Screenshot of an example summary report generated during the subtype discovery workflow.

### Figure S6

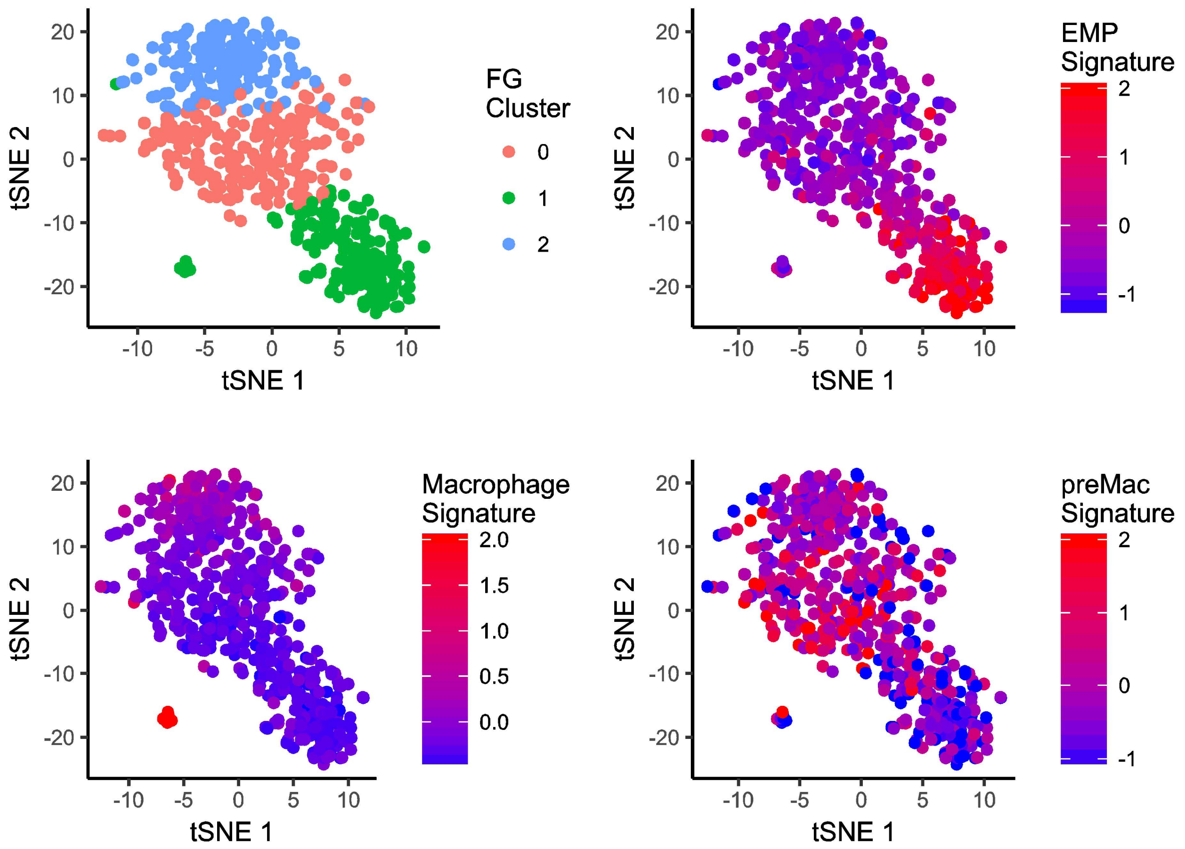


**Supplementary Figure 6: Re-Analysis of Mass et al. (A)** Cluster assignments of individual cells. **(B)** Individuals cells colored according to EMP signature gene expression. **(C)** Individual cells colored according to macrophage signature gene expression. **(D)** Individual cells colored according to preMac signature gene expression.

| Name | Functionality | Algorithm/Method | Status |
|---|---|---|---|
| calc_batch_effect_classifier | Batch effect quantification | Random Forest | Active |
| calc_clustering_hdbscan | Cell clustering | HDBSCAN | Active |
| calc_clustering_louvain | Cell clustering | Louvain | Under Development |
| calc_count_normalize | Normalization | various | Under Development |
| calc_de_genes_diffrank | Detection of differentially expressed genes | Diffrank (Scanpy) | Active |
| calc_de_genes_glm | Detection of differentially expressed genes | Generalized Linear Model | Active |
| calc_de_genes_nonparametric | Detection of differentially expressed genes | Mann-Whitney U test | Active |
| calc_diffusion_pseudotime | Pseudo-temporal ordering of cells | Scanpy | Active |
| calc_dimreduction_autoencoder | Dimensionality reduction | Neural network | Active |
| calc_dimreduction_tsne | Dimensionality reduction | tSNE | Active |
| calc_filter_quality | Quality control | Detection rate filtering | Under Development |
| calc_functional_analysis | Enrichment analysis | Fisher's Exact test | Active |
| calc_list_filtering | Blacklist/whitelist filtering | ID list filtering | Under Development |
| calc_logreg_confusion | | Logistic regression | Active |
| calc_normalize_tfidf | Normalization | TF/IDF | Active |
| calc_tsvd | Dimensionality reduction | Single value decomposition | Active |

**Supplementary Table 1A:** Calculation Apps in FASTGenomics. The table lists the calculation apps available in FASTGenomics and currently under development. For each app, the name, functionality and used algorithm or method is specified.

| Name | Usage | Status |
| --- | --- | --- |
| viz_barchart | Data Quality | Active |
| viz_batch_effect | Data Quality | Active |
| viz_dataquality | Data Quality | Active |
| viz_confusionmatrix | Cluster Inference | Active |
| viz_heatmap | DE Genes, Functional Analysis | Active |
| viz_scatterplot | Clustering, Diffusion Pseudo-time | Active |
| viz_table | DE Genes | Active |
| viz_linechart | Gene dynamics | Under development/ soon in FAST Genomics |

**Supplementary Table 1B:** Visualization Apps in FASTGenomics. The table lists the visualization apps available in FASTGenomics and currently under development.

| ID | GEO Accession Code | Organism | Number of Genes | Number of Cells |
|---|---|---|---|---|
| Macosko et al., Cell (2015) | GSE63473 | Mouse | 21,605 | 49,300 |
| Mass et al., Science (2016) | GSE81774 | Mouse | 8,553 | 408 |
| Moignard et al., Nature Biotechnology (2015) | --- | Mouse | 46 | 3,934 |
| Nestorowa et al., Blood (2016) | GSE81682 | Mouse | 23,357 | 1,920 |
| Paul et al., Cell (2015) | GSE72857 | Mouse | 19,362 | 10,368 |
| Tirosh et al., Nature (2016) | GSE70630 | Human | 22,338 | 4,347 |
| Tirosh et al., Science (2016) | GSE72056 | Human | 22,333 | 4,645 |
| Zeisel et al., Science (2015) | GSE60361 | Mouse | 18,920 | 3,005 |
| Zheng et al., Nature Communications (2017) | --- | Human | 21,253 | 68,579 |
| Ziegenhain et al., Molecular Cell (2017) | GSE75790 | Mouse | 22,701 | 482 |

**Supplementary Table 2:** Description of datasets available in the FASTGenomics pipeline.
